# Extensive location bias of the GPCR-dependent translatome via site-selective activation of mTOR

**DOI:** 10.1101/2024.06.17.599400

**Authors:** Matthew J Klauer, Caitlin AD Jagla, Nikoleta G Tsvetanova

## Abstract

G protein-coupled receptors (GPCRs) modulate various physiological functions by re-wiring cellular gene expression in response to extracellular signals. Control of gene expression by GPCRs has been studied almost exclusively at the transcriptional level, neglecting an extensive amount of regulation that takes place translationally. Hence, little is known about the nature and mechanisms of gene-specific post-transcriptional regulation downstream of receptor activation. Here, we apply an unbiased multiomics approach to delineate an extensive translational regulatory program initiated by the prototypical beta2-adrenergic receptor (β2-AR) and provide mechanistic insights into how these processes are orchestrated. Using ribosome profiling (Ribo-seq), we identify nearly 120 novel gene targets of adrenergic receptor activity which expression is exclusively regulated at the level of translation. We next show that all translational changes are induced selectively by endosomal β2-ARs. We further report that this proceeds through activation of the mammalian target of rapamycin (mTOR) pathway. Specifically, within the set of translational GPCR targets we discover significant enrichment of genes with 5’ terminal oligopyrimidine (TOP) motifs, a gene class classically known to be translationally regulated by mTOR. We then demonstrate that endosomal β2-ARs are required for mTOR activation and subsequent mTOR-dependent TOP mRNA translation. Together, this comprehensive analysis of drug-induced translational regulation establishes a critical role for location-biased GPCR signaling in fine-tuning the cellular protein landscape.

## INTRODUCTION

G protein-coupled receptors (GPCRs) are pivotal detectors and transducers of extracellular signals. At the cellular level, GPCRs respond to a range of sensory and chemical stimuli through induction of complex signaling networks that alter cell states. These begin with receptor-induced activation of heterotrimeric G proteins and proceed through the generation of second messengers, activation of effector kinases, and regulation of transcription and translation factors to ultimately re-wire gene expression in a stimulus-dependent manner.

Much is known about how GPCR signaling tunes gene transcription via biochemical regulation of select transcription factors (Ye 2001; Ho et al. 2009). Remarkably, it has recently emerged that these cellular responses are also regulated by GPCR localization, whereby the same receptor/G protein complex can elicit distinct responses depending on its subcellular site of signaling. In fact, the activation of transcription factors and regulation of gene transcription were among the first GPCR processes recognized as ‘location-biased’. Specifically, several receptors have been shown to induce transcriptional signaling from endosomal compartments, as opposed to the plasma membrane (Tsvetanova and Von Zastrow 2014; Godbole et al. 2017; Eiger et al. 2022). However, transcription is only the first of many steps that shape a cell’s protein repertoire in response to the environment. A significant amount of gene regulation takes place post-transcriptionally, and dedicated studies and global analyses have demonstrated that transcriptional and post-transcriptional regulatory programs often function as independent regulatory modules (Tebaldi et al. 2012; Moritz et al. 2019). Among many notable examples, activity-dependent translation of pre-existing mRNAs is a key mechanism underlying functional changes in the neuronal synapse linked to plasticity (Pfeiffer and Huber 2006). This underscores the importance of investigating each stage of gene regulation separately. Yet, the nature and mechanisms of gene-specific post-transcriptional regulation downstream of GPCR activation, and how these may be shaped by site-selective receptor activity, are poorly understood.

Here we leverage ribosome profiling (Ribo-seq) to define the GPCR-induced translatome and provide mechanistic insights into how these processes are orchestrated for a well-characterized receptor/second messenger pathway. We focus on the beta2-adrenergic receptor (β2-AR), a prototypical GPCR that mediates the effects of adrenaline and noradrenaline in the heart, lung and central nervous system via Gαs-dependent production of cyclic AMP (cAMP) (Madamanchi 2007; Mutlu and Factor 2008; O’Dell et al. 2015). We delineate an extensive novel β2-AR-dependent translational program and demonstrate that it is driven through spatially encoded crosstalk between the receptor and the mTOR pathway.

## RESULTS

### Ribosome profiling accurately captures actively translating RNA

We turned to a deep sequencing-based experimental platform that uncouples the contributions of transcription and translation to gene expression regulation. Specifically, we utilized a multiomics approach that integrates conventional RNA sequencing to measure mRNA expression (RNA-seq, “transcriptome”), with ribosome profiling to quantify isolated ribosome-protected fragments of RNA (RPFs) and define ribosome positions across each transcript at nucleotide resolution (Ribo-seq, “translatome”) **(Fig. 1a)**. While conventional proteomics methods measure steady-state protein levels, which reflect regulation at the stages of translation and protein stability, Ribo-seq is a powerful method to specifically interrogate genome-wide mRNA translational status that is not confounded by protein stability effects.

**Figure 1:**
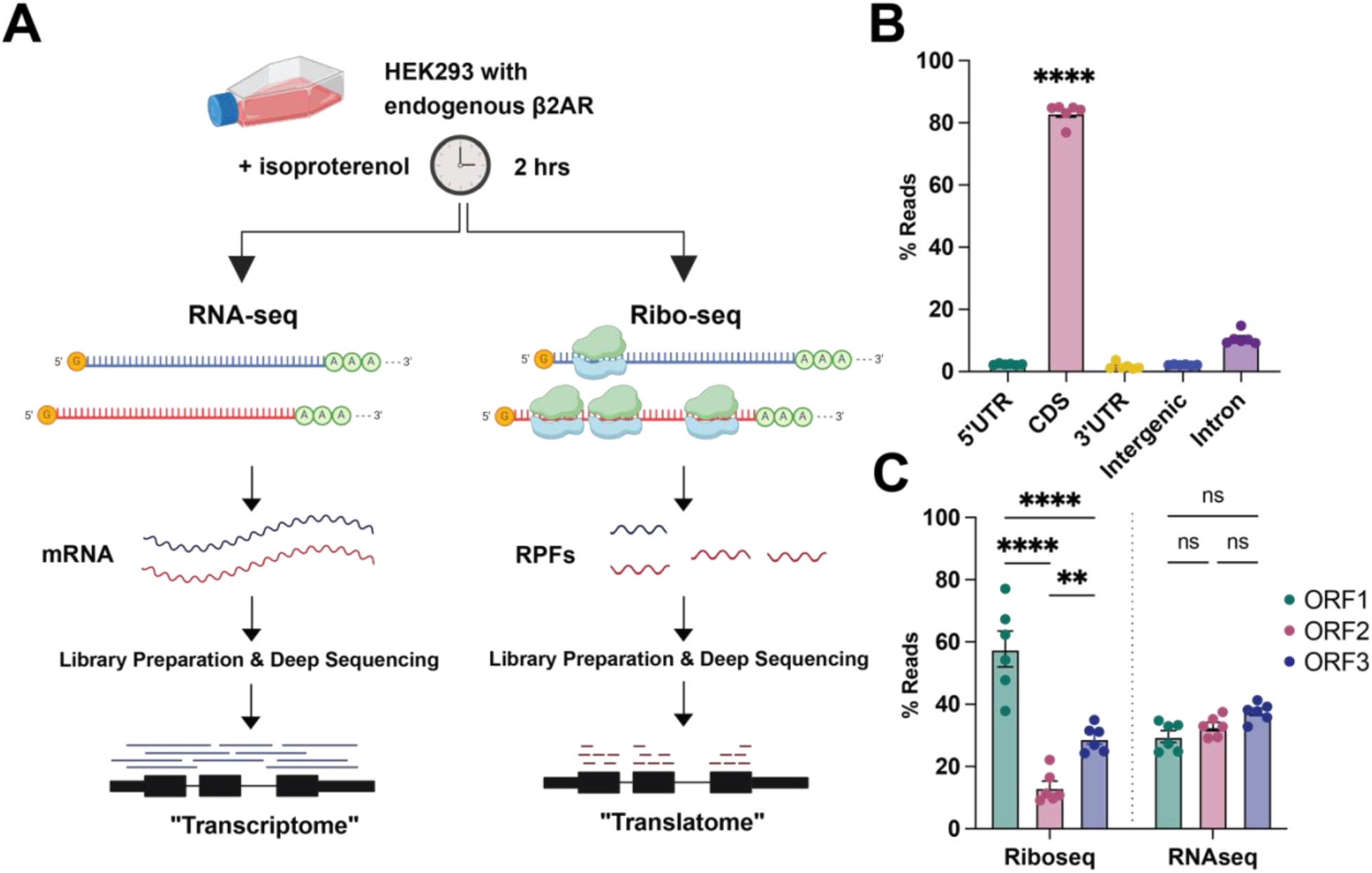
A snapshot of active global translation downstream of endogenous GPCR activity. (A) Schematic of the multiomics approach. HEK293 cells were treated with 1 µM Isoproterenol for 2 h. Lysates were prepared in parallel for sequencing of total RNA (“mRNA”, RNA-seq) and ribosome-protected fragments (“RPFs”, Ribo-seq). (B) Ribo-seq reads were mapped to genomic feature type using Ribotoolkit (Liu et al. 2020). (C) Ribo-seq and RNA-seq reads of RPF length (28-30 nt) evaluated for three nucleotide periodicity using Ribo-TISH (Zhang et al. 2017). Data are mean of n = 6 biological replicates. Error bars = ± s.e.m. **** = *p* < 0.0001, ** = *p* < 0.01 by one-way ANOVA test with Dunnett (B) or two-way ANOVA test with Tukey (C).

To examine these regulatory processes under native conditions, we selected HEK293 cells as a model, since these express endogenous β2-ARs (Violin et al. 2008). We stimulated adrenergic receptor signaling with the synthetic agonist isoproterenol for 2 hours, and then split each lysate for parallel analysis by RNA-seq and Ribo-seq **(Fig. 1a)**. We observed almost perfect correlation between biological replicates within each experimental setup, indicating high degree of reproducibility (Pearson coefficient > 0.95, *p* < 2.2 x 10^-16^, **Supplementary Fig. S1a-b)**. At the same time, replicates across the two platforms had Pearson coefficients of 0.40-0.50 (RNA-seq versus Ribo-seq, **Supplementary Fig. S1c)**. This confirms that each method is reporting a distinct layer of gene expression regulation. Further, the Ribo-seq experiments successfully captured ribosome-occupied mRNAs based on several quality control metrics. First, we observed enrichment of fragments of characteristic RPF length within coding sequences (CDS) relative to untranslated regions (UTRs) **(Fig. 1b, Supplementary Fig. S1d)**. Second, the aligned positions of RPFs revealed the expected stereotypical triplet periodicity within CDS, representing translocating ribosomes by three-nucleotide codon **(Fig. 1c)**. Notably, these trends were neither expected nor observed in the RNA-seq data **(Fig. 1c, Supplementary Fig. S1e-f)**.

Given the high degree of reproducibility and the strong enrichment for known translational features observed in the Ribo-seq experiments, we next set out to identify target genes with significant change in translational status in response to β2-AR activation.

### β2-AR activation mediates distinct transcriptional and translational programs

To define the β2-AR-dependent translatome, we independently identified “transcriptional” and “translational” target genes based on differential expression analysis of the untreated and isoproterenol-stimulated conditions in the RNA-seq and Ribo-seq data, respectively.

Starting with the RNA-seq dataset, we detected 61 genes with isoproterenol-induced changes in mRNA abundance (**Supplementary Table 1**). Among these, a significant number were already established as β2-AR transcriptional targets. Specifically, 17/61 genes were previously reported targets of β2-AR signaling by DNA microarray analysis of the same cell line under identical induction conditions (*p* < 2.2 x 10^-16^ by Fisher’s exact test). Next, we assessed agonist-stimulated changes in RPF abundance that did not exhibit concordant changes in RNA to pinpoint genes that are subject to translational regulation. For that, we used the change in the ratio of ribosome footprint to mRNA abundance within each gene’s CDS, a metric referred to as differential translation efficiency (ΔTE). We found 133 genes with ΔTE values of at least 1.5 standard deviations above or below the mean that were classified as translationally regulated β2-AR targets **(Fig. 2a, Supplementary Table 2)**. From these, we further distinguished two sub-categories of translational targets: “buffered” and “exclusive”. Specifically, we identified 15 translationally “buffered” genes that exhibited measurable transcriptional and translational changes in response to agonist, but only the transcriptional changes were statistically significant **(Fig. 2a**, light blue). On the other hand, the ΔTE for the remaining 118 genes was driven solely by change in RPF abundance with no significant transcription, and therefore these targets were classified as “exclusive” **(Fig. 2a-b**, red). For all further analyses, we focused on the set of exclusive targets as these represent extensively translationally regulated events.

**Figure 2:**
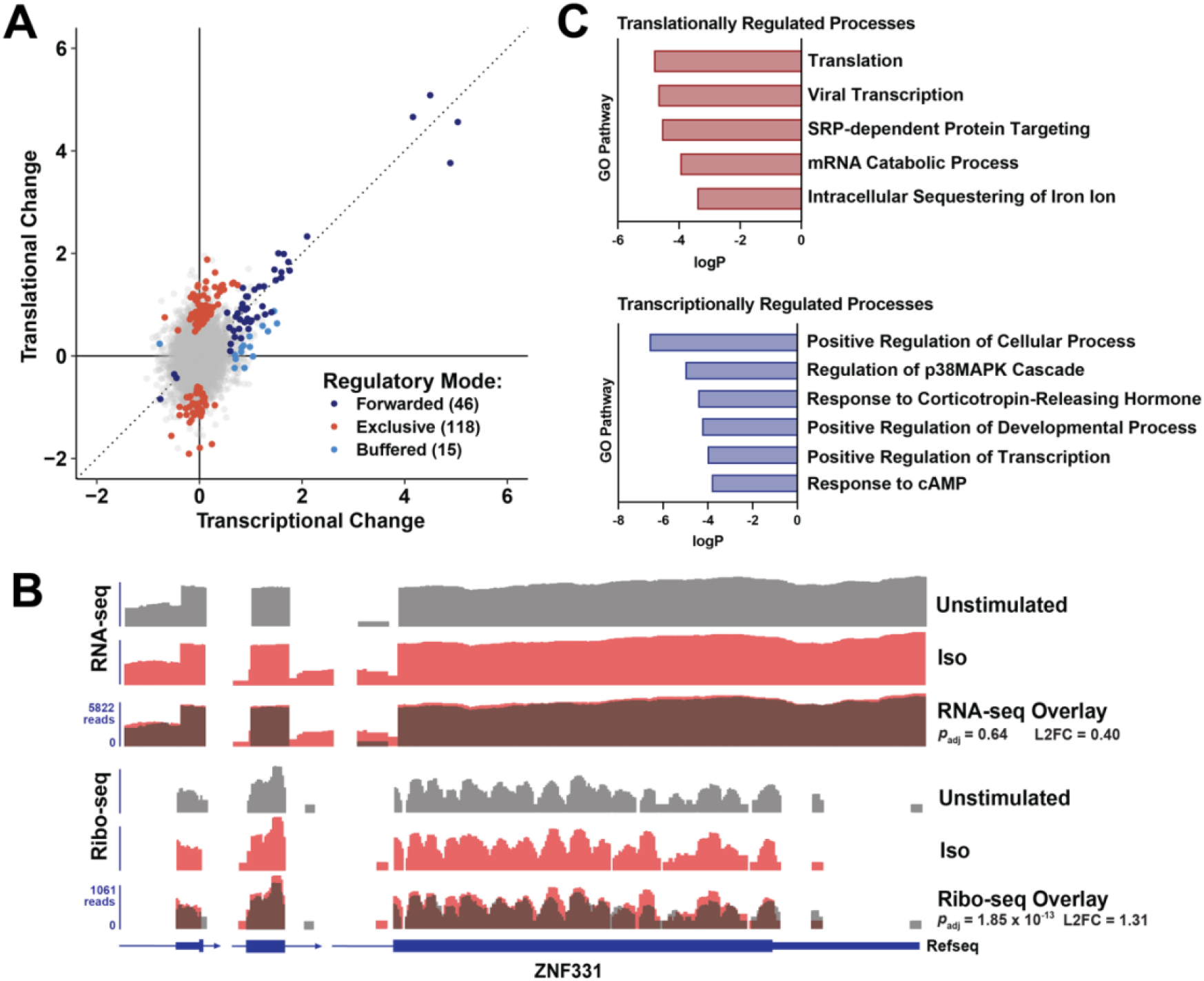
Comparison of transcriptional and translational regulation of gene expression in response to GPCR stimulation. (A) Scatter plot depicting changes in mRNA and RPF abundance (log_2_FC) following 2 h treatment with 1 µM Isoproterenol (“Iso”). Hits were categorized by significance and ΔTE as described in “Materials and Methods”. (B) IGV browser tracks showing normalized read coverage for select exons (boxes) and introns (lines) for the “exclusive” target, *ZNF331*. Iso treatment leads to an increase in ZNF331 translation with no change in mRNA abundance. The listed *p*_*adj*_ values were determined by Wald test in DeSeq and log2-fold change (L2FC) were calculated between Iso-treated and unstimulated samples (n=3 biological replicates). (C) Enriched Gene Ontology categories for “exclusive” translationally regulated genes (red, top) and transcriptionally regulated genes (blue, bottom). FDR < 0.05 was used as cut-off. All underlying data are summarized in **Supplementary Tables 1-3**.

Consistent with an autonomous nature of each gene regulatory mode, we noted several aspects that set apart the β2-AR translational from the transcriptional responses. First, the number of translational targets greatly exceeded the transcriptional repertoire, further underscoring translation as a key regulatory response downstream of GPCRs. Second, while the expression of virtually all transcriptional targets was increased by β2-AR signaling, the translational targets featured both induced and repressed genes (∼70 and ∼40 genes, respectively; **Supplementary Table 2)**. Presumably, the induction of GPCR-dependent transcriptional was mediated by CREB (cAMP response element-binding protein) (Altarejos and Montminy 2011). Supporting this established mechanism, our RNA-seq set contained a very significant number of CREB target genes (47/61 genes = 77%, *p* < 1.0 x 10^-19^ by Fisher’s exact test). In the case of translation, multiple regulatory pathways may exist allowing for both up- and down-regulation of genes downstream of the receptor. Lastly, to provide additional insights into the functions of each gene set, we performed Gene Ontology (GO) analysis. Comparison of the enriched GO categories revealed distinct biological functions associated with each regulatory mode **(Fig. 2c)**. Downstream of β2-AR stimulation, the induction of genes encoding factors with roles in cell differentiation, RNA polymerase II-dependent transcription, response to hormone stimulus, and MAPK regulation was mediated transcriptionally. On the other hand, genes encoding factors involved in translation, protein targeting, mRNA catabolism, and iron homeostasis were regulated at the level of translation **(Fig. 2c, Supplementary Table 3)**. Furthermore, network analysis of each GO category found that the encoded proteins participate in shared complexes, suggesting translational co-regulation among associating proteins **(Supplementary Fig. S2a-b)**. Therefore, β2-AR signaling gives rise to discrete transcriptional and translational programs with unique cellular processes regulated by each mechanism.

### Translationally regulated genes are mediated by endosomal β2-ARs

The mechanisms that mediate translation downstream of GPCR activation are poorly understood. Following ligand binding, the β2-AR stimulates Gαs-dependent production of cAMP first from the plasma membrane and subsequently from early endosomes upon internalization (Tsvetanova and Von Zastrow 2014; Irannejad et al. 2017). Therefore, as a first step in disentangling how β2-ARs regulate translation, we investigated whether and how the response may be shaped by receptor localization.

To confine β2-AR activation to the plasma membrane, we used an established pharmacological method to acutely inhibit dynamin-dependent endocytosis with the drug Dyngo-4a (“Dyngo”) (Harper et al. 2011; Tsvetanova and Von Zastrow 2014; Irannejad et al. 2017) **(Supplementary Fig. S3a)**. Then, we examined gene expression changes by parallel RNA-seq and Ribo-seq analyses **(Supplementary Fig. S3b)**. Comparison between the unstimulated conditions revealed that Dyngo did not impact basal mRNA or RPF abundance, supporting that inhibitor treatment alone does not alter steady-state transcription or translation **(Supplementary Fig. S3c-d)**. We previously established that β2-AR internalization is required for efficient activation of the downstream transcriptional responses (Tsvetanova and Von Zastrow 2014). In agreement, the induction of transcriptional targets identified here through RNA-seq was significantly blunted by Dyngo (*p* < 1.1 x 10^-10^ by Wilcoxon test, **Supplementary Fig. S3e)**. This supports that the approach is effective at recapitulating known biology. Remarkably, we observed that blockade of endocytosis also led to pervasive inhibition of the β2-AR-dependent translational regulation (*p* < 1.7 x 10^-13^ by Wilcoxon test, **Fig. 3)**. Strikingly, we found an even greater dependence of gene translation on internalized receptors compared to gene transcription. Specifically, none of the 118 translational targets identified in cells with normal endocytosis undergo significant changes in translation following endocytic blockade (using *p* < 0.05 cut-off by Wald test comparing untreated and isoproterenol-stimulated conditions in the presence of Dyngo). On the other hand, more than a fifth of the β2-AR transcriptional targets (14/61 genes = 23%) were mildly but significantly induced by isoproterenol also in the presence of Dyngo **(Supplementary Table 1)**. Hence, gene translation is regulated exclusively by internalized β2-ARs.

**Figure 3:**
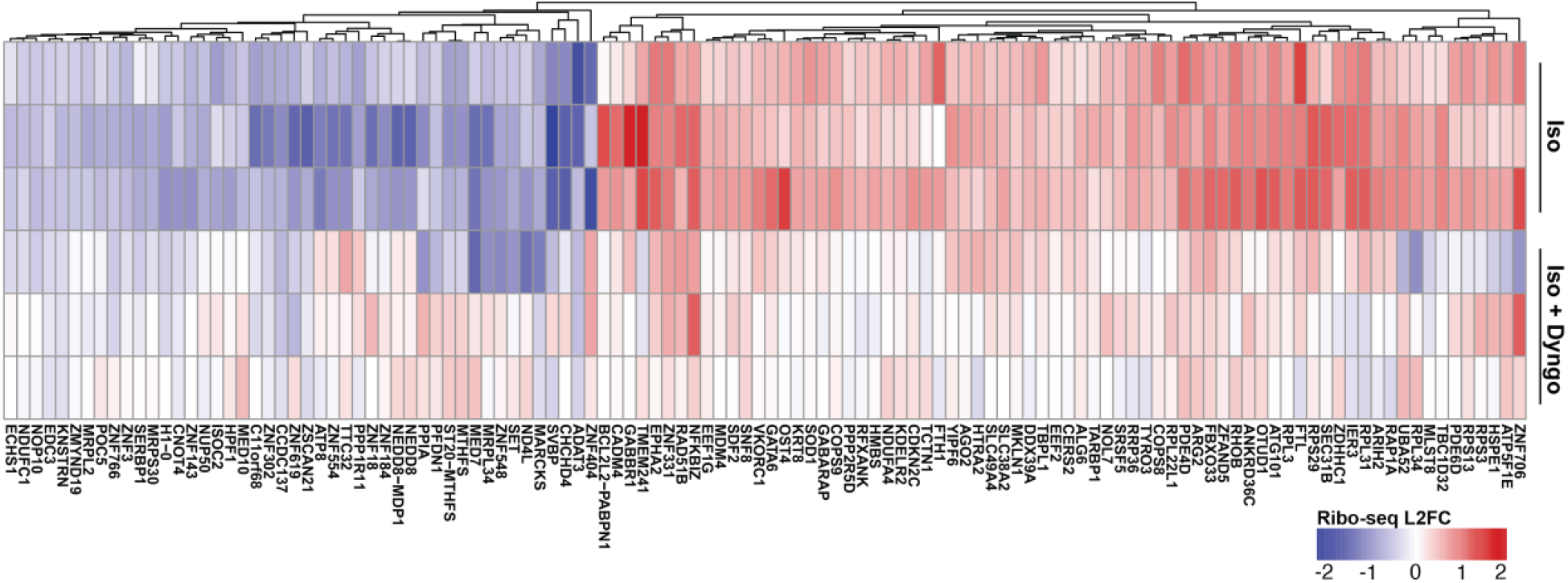
The GPCR-dependent translatome is spatially encoded. (A) Heatmap of the exclusive translational targets. Colors represent Log2 fold change (L2FC) between Isoproterenol (“Iso”)-treated vs unstimulated RPFs in the presence of vehicle (DMSO) or 30 µM Dyngo. K-means clustering was performed with the R-package pheatmap.

### Endosomal β2-ARs activate mTOR to modulate TOP mRNA translation

To further pinpoint how endosomal β2-AR signaling mediates protein synthesis, we focused on categories of genes with robust isoproterenol-dependent changes in our Ribo-seq analysis. We recognized that the translation efficiency of mRNAs containing annotated terminal oligopyrimidine (TOP) motifs was disproportionally induced by receptor stimulation (*p* < 7.7 x 10^-3^ by Kolmogorov– Smirnov test, **Fig. 4a**), and a significant number of TOP mRNAs were among the set of ∼120 robust translational targets (*p* < 1.0 x 10^-10^ by Fisher’s exact test). Importantly, this regulation required intact endocytosis (*p* < 1.6 x 10^-3^ by Wilcoxon test). TOP mRNAs primarily encode components of the translation machinery and nearly all ribosomal proteins (Thoreen et al. 2012). Consistent with this, “Translation” was the most significantly impacted Ribo-seq GO category (*p* < 1.5 x 10^-5^, **Fig. 2c**). Mechanistically, the synthesis of ribosomal proteins, including TOP mRNAs, is regulated by the mTOR pathway. Therefore, we surmised that endosomal β2-ARs may selectively activate mTOR signaling to modulate the translation of this gene class.

**Figure 4:**
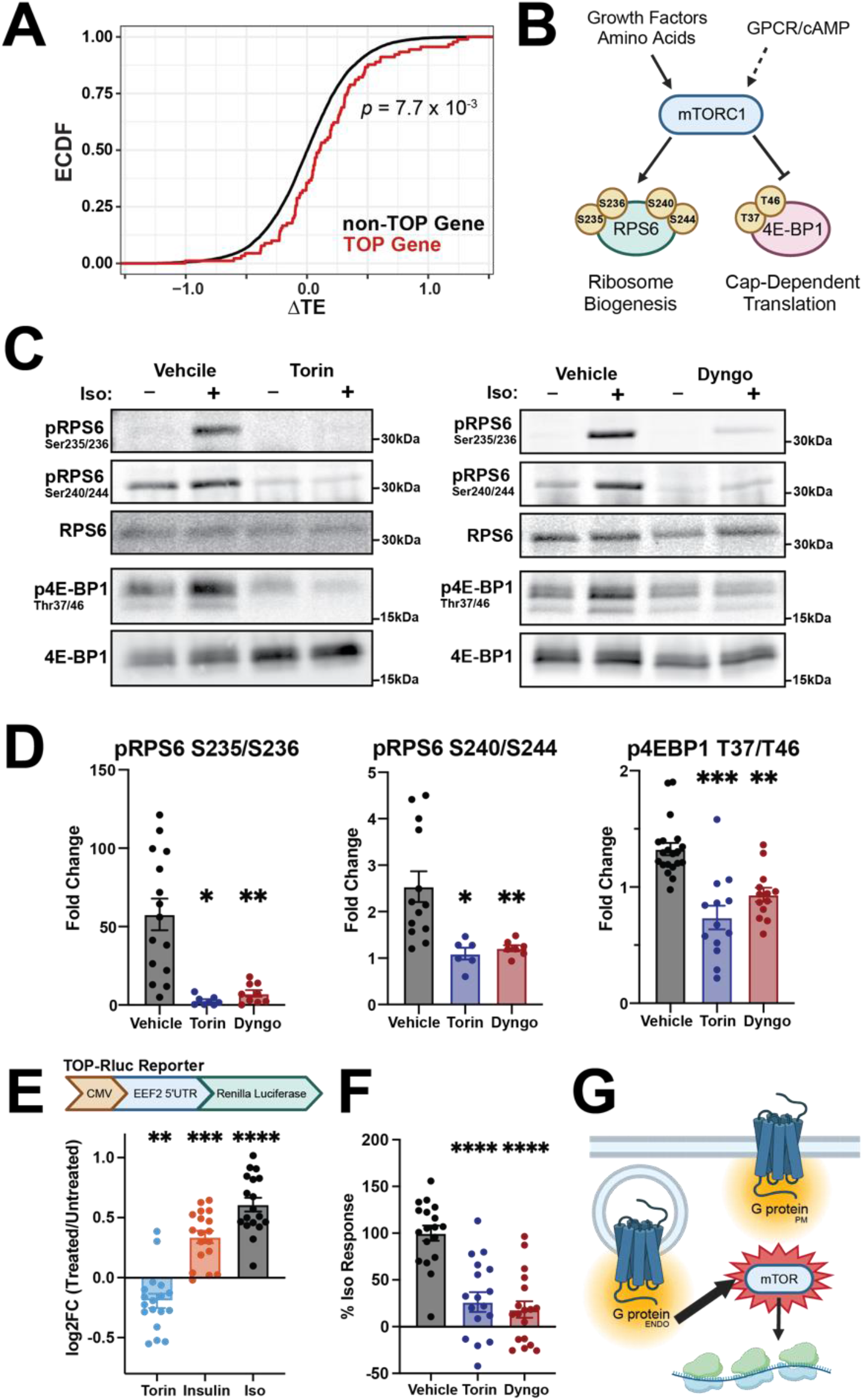
Endosomal β2-AR regulate TOP mRNA translation via mTOR activation. (A) Empirical cumulative distribution function (ECDF) of the differential translation efficiency for all genes in response to stimulation with 1 µM Isoproterenol (“Iso”). The distribution of genes with TOP motifs is shown in red. The resultant *p*-value was derived by Kolmogorov-Smirnov test comparing the distributions of TOP and non-TOP motif-containing genes. (B) Schematic of mTOR-dependent regulation of translation via phosphorylation of RPS6 and 4E-BP1. (C-D) Endosomal β2-AR signaling is required for mTOR-dependent phosphorylation of RPS6 and 4E-BP1. HEK293 cells were grown in serum-free medium for 24 h, then pre-treated with vehicle (DMSO), 100 nM Torin, or 30 µM Dyngo-4a for 20 min. Cells were then stimulated with 1 µM Iso for 10 min and lysates were subjected to Western blot analysis with antibodies against total and phosphorylated RPS6 and 4E-BP1. Fold-change values represent the ratio of phosphorylated/total protein in Iso-treated vs unstimulated conditions. All data are mean of n=6-20 biological replicates. (E) HEK293T cells were transfected with TOP-Rluc reporter (schematic, top) and a control plasmid encoding firefly luciferase (control-Fluc) for 24 h, then treated with 100 nM Torin, 100 nM Insulin, or 1 µM Iso for 6 h. Cells were measured by dual-luciferase assay and reporter translational induction was calculated as Rluc/Fluc and shown as L2FC (stimulated vs unstimulated cells) (bottom). (F) Endosomal β2-AR signaling activates TOP-mRNA translation via mTOR. HEK293T cells were transfected as in (E), pre-treated with vehicle (DMSO), 100 nM Torin, or 30 µM Dyngo for 20 min, then stimulated with 1 µM Iso for 6 h. Data are shown as percent of the maximum response measured with Iso in the presence of vehicle. All data are mean of n=18 biological replicates. (G) Model: Endosomal β2-AR signaling is required for TOP gene translation via site-selective activation of mTOR. Error bars = ± s.e.m. **** = *p* < 0.0001, *** = *p* < 0.001, ** = *p* < 0.01, * = *p* < 0.05 by unpaired Student’s *t*-test comparing vehicle to inhibitor-treated conditions in (D) and one-way ANOVA test with Dunnett in (E-F).

To test this, we began by interrogating whether β2-AR signaling impacts mTOR activity. Eukaryotic Initiation Factor 4E-Binding Protein 1 (4E-BP1) is phosphorylated directly by mTOR on residues Thr37/46, and ribosomal protein 6 (RPS6) is phosphorylated on sites Ser235/236 and Ser240/244 (Battaglioni et al. 2022; Meyuhas 2015) **(Fig. 4b)**. Therefore, we monitored accumulation of phosphorylated 4E-BP1 and RPS6 by Western blot analysis as markers of mTOR activation. Acute stimulation of β2-AR with isoproterenol led to significant induction of 4E-BP1 and RPS6 phosphorylation **(Fig. 4c-d**, black bars). These events were dependent on mTOR, as pre-treatment with the ATP-competitive inhibitor, Torin (Thoreen et al. 2009), abolished the regulation **(Fig. 4c-d**, blue bars). The mTOR response also required PKA activity downstream of the β2-AR, consistent with a necessity for cAMP in this process **(Supplementary Fig. S4a)**. More importantly, acute application of Dyngo completely blocked mTOR-induced phosphorylation of 4E-BP1 and RPS6 in response to GPCR agonist **(Fig. 4c-d**, red bars). This supports that β2-AR endocytosis is essential to activate the mTOR pathway.

Next, we aimed to establish a connection between mTOR induction by intracellular receptors and the regulation of TOP gene translation. *EEF2* was among the TOP mRNAs significantly upregulated by β2-AR signaling based on the Ribo-seq analysis **(Supplementary Table 2)**. Therefore, as an orthogonal approach to monitor TOP mRNA translation, we obtained a validated reporter which utilizes Renilla luciferase expression downstream of the *EEF2* 5’UTR harboring its TOP sequence (“TOP-Rluc”) (Philippe et al. 2020). As expected, treatment with Torin alone led to TOP-Rluc repression, while insulin yielded a significant increase **(Fig. 4e)**. These represent responses to inhibition and activation of mTOR that are independent of GPCR signaling. Next, we assessed reporter levels under β2-AR induction conditions. TOP-Rluc expression was induced by isoproterenol in a PKA-dependent manner, and the magnitude of this change was greater than that following mTOR activation with insulin **(Fig. 4e, Supplementary Fig. S4b)**. These reflect reporter translation, because neither isoproterenol nor insulin impacted TOP-Rluc mRNA levels **(Supplementary Fig. S4c)**. We then asked whether mTOR signaling is required for this regulation. We measured reporter accumulation following β2-AR stimulation in the presence of an mTOR inhibitor. Co-application of Torin drastically repressed the isoproterenol-dependent accumulation of TOP-Rluc **(Fig. 4f**, blue bar). In support of the regulatory significance of the TOP motif, an analogous luciferase reporter which harbors a mutated TOP motif within its UTR (Philippe et al. 2020) displayed minimal β2-AR-dependent accumulation that was not affected by Torin (“TOPmut-Rluc”, **Supplementary Fig. S4d**). Lastly, endocytic blockade completely precluded the translational induction of TOP-Rluc in isoproterenol-stimulated cells, corroborating a significant role for receptor internalization in this process **(Fig. 4f**, red bar).

Collectively, these experiments delineate that the induction of TOP mRNA translation downstream of β2-ARs requires endocytosis and proceeds through an mTOR-dependent mechanism **(Fig 4g)**.

## DISCUSSION

Numerous extracellular signals affect physiological responses by binding to GPCRs, leading to changes in gene expression. However, understanding of how GPCRs influence the capacity of cells to dynamically regulate protein content in response to stimuli has remained limited, as studies on receptor-mediated gene expression have focused almost exclusively on transcription. The present work takes advantage of the sensitivity and high-throughput nature of Ribo-seq to delineate an extensive new regulatory program orchestrated by a GPCR that consists of nearly 120 genes whose translation is regulated independently of mRNA levels. These genes have not been previously associated with β2-AR and cAMP signaling, making them the first examples of drug-induced translational regulation mediated by adrenergic receptor activation. Notably, at a matched time point the repertoire of translationally regulated genes exceeds the transcriptional one, and each set orchestrates distinct biological functions and processes (**Fig. 2, Supplementary Table 3**). Therefore, by focusing on gene translation and its regulation by receptor activity, these findings provide novel insights into the comprehensive mechanisms by which GPCR signaling shapes cell states.

Over the past decade, it has been recognized that GPCR signaling is subject to intricate spatial regulation, and that the selective modulation of localized receptors can have distinct effects on physiology (Klauer, Willette, and Tsvetanova 2024). Despite this growing appreciation for the importance of compartmentalized GPCR signaling, how different subcellular fractions of receptors shape unique cellular behaviors has remained poorly understood. Our findings indicate that a salient molecular mechanism involves the exclusive coupling between gene translation and endosomal receptors **(Fig. 3)**. We further define how intracellular β2-ARs mediate these processes by demonstrating that the translational induction of TOP mRNAs by agonist proceeds through receptor endocytosis-dependent activation of mTOR **(Fig. 4)**. Of note, an interplay between the GPCR/cAMP and mTOR signaling pathways has been documented previously. Interestingly, the reported impact of receptor signaling on mTOR varies depending on the study, with some demonstrating that cAMP stimulates mTOR (Blancquaert et al. 2010; Liu et al. 2016) while others support inhibition (Xie et al. 2011; Jewell et al. 2019). Typically, these discrepancies have been attributed to differences in experimental models and/or experimental design (Shi and Collins 2023). However, the significance of localized receptor activity in the regulation of mTOR was not considered. Our findings ascertain signal compartmentalization as a new pertinent axis of regulation underlying the crosstalk between these two pathways. In fact, this is the first demonstration to our knowledge that crosstalk between GPCRs and any other signaling pathway is reliant on signal compartmentalization. Hence, we speculate that this unappreciated location-biased nature of the β2-AR/mTOR interplay could be a critical parameter to consider in dissecting the conflicting observations regarding the effects of cAMP on mTOR.

An unresolved puzzle arising from our discoveries is how GPCR activity and its spatial bias modulate the mTOR pathway. Despite the documented examples of interplay between the cascades, a commonly agreed-upon detailed mechanistic model linking GPCRs and mTOR is mostly lacking. Various groups have shown that the regulation requires PKA (Liu et al. 2016; Blancquaert et al. 2010; Jewell et al. 2019), which is consistent with our data **(Supplementary Fig. S4**). Further, previous reports have suggested that the GPCR effects are independent of several known mTOR regulatory inputs, including Rheb, the Rheb GAPs TSC1/2, AMPK, or Rag GTPases (Xie et al. 2011; Jewell et al. 2019). Instead, the regulation may proceed through direct PKA-dependent phosphorylation of mTOR complex 1 (mTORC1) (Liu et al. 2016; Blancquaert et al. 2010; Zhang et al. 2021), although it is not clear whether and which of the documented events take place across multiple contexts, and which are context-specific. For example, it was shown that PKA phosphorylates the mTOR kinase itself in one model (Xie et al. 2011) but not in another (Jewell et al. 2019). Similarly, while PKA can phosphorylate the mTORC1 negative regulator PRAS40 in vitro (Blancquaert et al. 2010), it remains to be determined whether this takes place in cells. One mechanism that has garnered independent support from multiple groups involves the phosphorylation of RAPTOR (Regulatory-Associated Protein of mTOR) by PKA. This process was found to be required in the context of both activation and inhibition of mTOR by cAMP (Xie et al. 2011; Jewell et al. 2019), although it remains to be determined how RAPTOR phosphorylation alters mTORC1 activity. Nevertheless, a recent report uncovered that this is dependent on the scaffolding protein, AKAP13 (A-kinase anchor protein 13), which brings PKA in physical proximity to mTORC1 (Zhang et al. 2021). AKAPs are of recognized significance in defining cAMP/PKA compartmentalization through the scaffolding of multimolecular “signalosomes” (Omar and Scott 2020). It is tempting to speculate that the AKAP13/PKA/RAPTOR complex may render that “nanodomain” of PKA selectively responsive to increase in local cAMP concentrations and thus play an important role in establishing the location bias of GPCR-dependent mTOR regulation. It is pertinent to examine this directly and to further dissect the precise mechanisms through additional candidate-based and unbiased approaches.

In summary, our study provides a comprehensive analysis of how GPCR signaling impacts gene translation and presents novel mechanistic insights into these processes. Ultimately, this work extends the concept of compartmentalized receptor activity to a previously underexplored layer of gene expression regulation to fundamentally enrich the current understanding of the remarkable diversity of endogenous cellular responses and their location bias. While we demonstrate mTOR as one of the downstream pathways through which intracellular receptors exert translational control, much remains to be learned about the precise molecular underpinnings of this interplay. The mTOR pathway is a master regulator of cell growth and proliferation that is impaired in many pathological conditions, including cancer, metabolic disorders and neurodegeneration (Laplante and Sabatini 2012). Given the equally critical functions of GPCRs as coordinators of human physiology and pathophysiology and their immense pharmacological potential (Sriram and Insel 2018), further mechanistic dissection of the GPCR/mTOR crosstalk may help identify novel candidates for therapeutic targeting in the context of human diseases arising from dysfunctions in these pathways. Along these lines, endosomal β2-ARs likely also mediate gene translation via mTOR-independent mechanisms. The elucidation of additional mechanisms linking GPCR signaling to gene translation is therefore another key avenue for future exploration. Lastly, we envision that these analyses can be extended to investigate how GPCR activation impacts global as well as local mRNA translation in other cell systems, including neurons, where protein synthesis from pre-existing mRNAs is of recognized physiological importance.

## ACKNOWLEDGEMENTS

We thank Dr. Carson Thoreen (Yale University, CT) for his valuable insights regarding mTOR regulation of TOP genes and for generously providing the TOP reporter constructs. We also thank Dr. Nicholas Ingolia (UC Berkeley, CA) for helpful discussions regarding Ribo-seq protocols. This work was supported by the National Institutes of Health (R01NS127847 to N.G.T.) and American Heart Association (19IPLOI34670002 to N.G.T). Figures 1a, 4b and 4g were created using BioRender.

## CONTRIBUTIONS

N.G.T. supervised the project. N.G.T., C.A.D.J. and M.J.K. conceived the project. C.A.D.J. carried out the initial experimental and bioinformatics Ribo-seq adaptations. M.J.K performed and analyzed all experiments included in this study. N.G.T. and M.J.K interpreted the results and wrote the manuscript.

## MATERIALS AND METHODS

### Chemicals and Antibodies

(-)-**Isoproterenol hydrochloride** (Sigma-Aldrich, Cat #I6504) was dissolved in water/100 mM ascorbic acid to 10 mM stock and used at 1 μM final concentration. **Dyngo-4a** (Abcam, Cat #120689) was resuspended in DMSO to 30 mM stock and used at 30 μM final concentration. **Torin-1** (MedChemExpress, Cat #HY-13003) was resuspended to 100 μM in DMSO and used at a final concentration of 100 nM. **Insulin** (Sigma -Aldrich, Cat #91077C) was resuspended to 1 mM in water/HCl (pH 2.0) and used at a final concentration of 100 nM. **Coelenterazine** (GoldBio, Cat #CZ) was resuspended to 10 mM in ethanol and stored protected from light. **D-luciferin** (Goldbio, Cat# LUCNA-100) was resuspended in 10 mM HEPES buffer and used at 150 μg/mL final concentration. **H89** (Cayman Chemical, Cat #10010556) was dissolved in DMSO to 10 mM and used at 10 μM final concentration.

**RPS6** mouse monoclonal antibody (Cell Signaling Technologies, Cat #2317), **phospho-RPS6** (S235/S236) rabbit monoclonal antibody (Cell Signaling Technologies, Cat #4858), **phospho-RPS6** (S240/S244) rabbit monoclonal antibody (Cell Signaling Technologies, Cat #5364), **4E-BP1** rabbit monoclonal antibody (Cell Signaling Technologies, Cat #9644), and **phospho-4E-BP1** (T37/T46) (Cell Signaling Technologies, Cat #2855) were all used at 1:1000 for Western blots. The following secondary antibodies were used at 1:10,000 dilutions: donkey anti-mouse-680 (LICOR Biosciences, Cat #926-68072) and donkey anti-rabbit-800 (LICOR Biosciences, Cat #926-32213).

### Cell Culture

HEK293 cells were obtained from ATCC and HEK293T cells were obtained from Clontech. Cell lines were grown at 37°C/5% CO2 in Dulbecco’s Modified Eagle Medium (Thermo Fisher Scientific, Cat #11965118) supplemented with 10% fetal bovine serum (Sigma-Aldrich, Cat #F2442). HEK293 cells with stable Flag-β2-AR for internalization assays were previously described (Semesta et al. 2020).

### Ribosome Profiling

Samples were harvested and ribosome footprints generated using the standard strategy of McGlincy and Ingolia (McGlincy and Ingolia 2017) with slight modifications to facilitate the use of commercial kits for rRNA depletion and sequencing library preparation. The monosome isolation step was eliminated, as previous work has shown that size selection for RNA fragments with lengths characteristic of RPFs (∼25-35 nt) is sufficient for capturing actively translating mRNA (Zappulo et al. 2017).

HEK293 cells were plated in 15 cm dishes for 70% confluency on the day of the experiment. For endosome inhibition experiments, cells were pre-incubated for 20 min in serum-free DMEM with either 30 µM Dyngo-4a or DMSO. Cells were stimulated with 1 µM isoproterenol in serum-free DMEM for 2 h, then rinsed twice with ice-cold PBS (GenClone, Cat #25-507) and lysed in-dish with 400 µL freshly prepared lysis buffer [20 mM Tris (Sigma, Cat #T2194), 150 mM NaCl (Thermo, Cat #AM9759), 5 mM MgCl2 (Thermo, Cat #AM9530G), 1 mM DTT (Sigma-Aldrich, Cat #43816), 1% Triton X-100 (Acros Organics, Cat #AC32737100), 100 µg/mL cycloheximide (Sigma-Aldrich, Cat #C4859), and 25 U/mL TURBO DNase (Thermo, Cat #AM2238)]. Following incubation in lysis buffer for at least 10 min on ice, lysates were triturated 10 times through a 25G needle then clarified by centrifugation (10 min/20,000 x g/4°C) prior to flash-freezing in liquid nitrogen and storage at -80°C. An aliquot of lysate for each sample was purified with the Zymo RNA Clean & Concentrator-25 kit (Genesee, Cat #11-353) following manufacturer’s protocol for later use in preparing poly-A selected mRNA libraries.

An aliquot of lysate containing 30 µg total RNA was taken for input to the ribosome profiling workflow. After bringing the volume to 200 µL with freshly prepared polysome buffer (lysis buffer without cycloheximide or DNase), lysates were incubated with 15 U (1.5 µL) RNase I (VWR, Cat #76081-704) for 45 min at room temperature with gentle agitation. The digestion reaction was terminated by adding 200U (10 µL) SUPERase RNase inhibitor (Thermo, Cat #AM2696) and 20 µL 10% SDS (Promega, Cat #V6551) and then purified using the Zymo RNA Clean & Concentrator kit (Genesee, Cat #11-353/11-325), following manufacturer’s modified protocol to isolate small RNAs (< 200 nt).

The entirety of the eluate was input to an end repair reaction for 1 h at 37°C to generate 3’-OH/5’-PO4 RNA fragments compatible with downstream adapter ligation. Final reaction volume of 40 µL contained ∼ 24 µL of sample, 40U (4 µL) T4 polynucleotide kinase enzyme (T4 PNK; NEB, Cat #M0201S), 4 µL 10X T4 Ligase Buffer (NEB, Cat #B0202S; final reaction concentrations of 50 mM Tris-HCl, 10 mM MgCl2, 1 mM ATP, 10 mM DTT, pH 7.5), 4 µL 10 mM ATP (NEB, Cat #P0756S), 40U (2 µL) SUPERase inhibitor, and nuclease-free water. Reactions were terminated by transferring the tube to ice, and samples were purified with the same modified protocol for the Zymo RNA Clean & Concentrator kit. A supersaturating final reaction concentration of 2 mM ATP was chosen because T4 PNK enzymatic activity is irreversible and therefore, flooding the reaction serves to minimize the impact of between-sample differences in digestion or total RNA quantity on availability of fragments compatible with downstream adapter ligation.

Ribosome-protected fragments (RPFs) between ∼25-35 nt were size-selected by gel electrophoresis on 15% TBE-Urea gels (Bio-Rad, Cat #4566055), then eluted via crush-and-soak extraction in buffer containing 300 mM NaOAc (Thermo, Cat #AM9740), 1 mM EDTA (Thermo, Cat #AM9260G), 10% SDS (Promega, Cat #V6551). Crushed RPF-containing bands were eluted overnight at 4°C, then slurries were centrifuged through Spin-X cellulose filters (Sigma-Aldrich, Cat #CLS8163) at 14,000 RPM for 2 min. To precipitate RNA from clarified eluates, 20 µL of 3M NaOAc and 2 µL Glycoblue (Thermo, Cat #AM9515) were added and mixed well, followed by addition of 500 µL isopropanol and further vigorous mixing before overnight precipitation at -80°C. Then, eluates were centrifuged for 1 hr at 21,100 x g at 4°C and pellets containing RPFs were rinsed once with ice-cold freshly diluted 80% ethanol, air dried by 5 min Speedvac at room temperature, and resuspended in 14 µL nuclease-free water.

Samples were depleted of rRNA using the Human riboPOOL for Ribosome Profiling RNA kit (Galen Molecular, Cat #dp-K024-000042) with a modified protocol omitting the 50°C/5 min incubation following magnetic bead addition to the hybridized RNA-rRNA oligo reaction. After aspirating supernatant away from rRNA-capturing magnetic beads, RPFs were further purified using the small RNA purification protocol for the Zymo RNA Clean & Concentrator kit and Zymo-5 columns and eluted in 6 µL nuclease-free water. The entire yield was used to generate sequencing libraries following manufacturer’s protocol for NEBNext Multiplex Small RNA Library Prep Set for Illumina (New England Biolabs, Cat #E7300S), using 12 PCR cycles. The final product was purified using the Zymo DNA Clean & Concentrator-5 kit (Genesee, Cat #11-302) following a modified protocol for small DNA fragments (< 200 nt), and ∼140-150nt DNA fragments were then size-selected by gel electrophoresis with 5% TBE gels (Bio-Rad, Cat #4565015) following NEB library prep kit protocol. Libraries were sequenced with Illumina HiSeq 4000, PE150.

### Polyadenylated mRNA Library Preparation

Total RNA was isolated from an aliquot of each lysate above using manufacturer’s standard protocol for the Zymo RNA Clean & Concentrator-25 kit (Genesee, Cat #11-353) and 1 µg total RNA was used as input to NEBNext Poly(A) mRNA Magnetic Isolation Module for NEBNext Ultra II RNA Library Prep Kit for Illumina (New England Biolabs, Cat #E7775), following the manufacturer’s protocol and using 7 PCR cycles for library amplification. Additional unpaired mRNA libraries were prepared using NEBNext Ultra II Directional RNA Library Prep Kit for Illumina, following the manufacturer’s protocol (New England Biolabs, Cat #E7776S). Libraries were sequenced on Illumina HiSeq 4000.

### RNA-seq and Ribo-seq Data Processing

Sequencing data were trimmed BBDuk (BBMap version #38.63) to remove adaptors. Sequences aligning to ncRNA, tRNA, and rRNA were removed using Bowtie (version 2.3.5) and cleaned files were aligned to the Genome Reference Consortium Human Build 38 (GRCh38) using STARaligner (version 2.7.5). FeatureCounts (Subread version 1.6.3) was used to align and quantify coding sequences, with RPF reads being narrowed to reads of length 26 to 34 nucleotides.

DESeq2 (version 3.16) was used for differential expression analysis of raw reads. Significant changes at either the RNAseq or Ribo-seq level were determined by Wald test with adjusted *p*-value of 0.05 (Love, Huber, and Anders 2014). For ΔTE calculations, unstimulated RNA samples were used as an additional covariate in the DESeq input as described in (Chothani et al. 2019). Genes with significant changes in mRNA or RPFs were further categorized based on their respective ΔTE values. “Exclusive” genes were defined as genes with ΔTE values at least 1.5 standard deviations above or below the mean in the same direction as the corresponding significant RPF change (*p* < 0.05 by Wald test). “Buffered” genes were defined as genes with ΔTE values less than 1.5 standard deviations below the mean in the same direction as the corresponding significant mRNA abundance change (*p* < 0.05 by Wald test). The remaining genes with significant mRNA changes were categorized as “Forwarded”.

Ribotish and Ribotoolkit were used for periodicity and feature-type analysis of the Ribo-seq data, respectively (Zhang et al. 2017; Liu et al. 2020). Feature count analysis of RNAseq data was performed using GenomicFeatures in R (Lawrence et al. 2013). Correlation analysis to compare Pearson correlation coefficients for log-transformed counts between conditions and across library preparations was performed using bigPint (Rutter and Cook 2020). For gene ontology analysis, categorized hits were entered into the GOrilla GO term analysis software and significantly regulated pathways were determined by FDR < 5%. Protein interaction matrices were generated by String-DB (Szklarczyk et al. 2023) and analyzed in Cytoscape (Shannon et al. 2003). Genomic alignment traces were generated in Interactive Genomics View (IGV) (Robinson et al. 2011) and modified in Adobe Illustrator.

### TOP-Rluc Reporter Assay

HEK293T cells were seeded on 6-well dishes for 24 h, then transfected with TOP-Rluc or TOPmut-Rluc and Firefly Luciferase plasmids using Lipofectamine-2000 (Invitrogen, Cat #11668027) following recommended protocols. After 24 h, cells were lifted with Accutase (Gibco, Cat #A1110501), pelleted, and resuspended in serum-free DMEM. Cells were split across black 96 well assay plates (Corning, Cat #3603) and incubated with inhibitor or DMSO for 20 min as described. Following 6 h stimulation with agonist, cells incubated with 150 μg/mL D-luciferin for 40 min and an initial Firefly luciferase reading was taken. Afterwards, cells were lysed in Renilla luciferase lysis buffer (4mM EDTA (Sigma, Cat #ED4SS), 1% glycerol (Sigma, Cat #G5516), 0.1% Triton X-100 (Sigma, Cat #X100), 50 mM HEPES, 1 mM DTT, and 8 µM coelenterazine). To measure luciferase signal, the plate was read on the Hamamatsu μCell plate reader at 37°C. Samples were normalized by diving Renilla luciferase by Firefly luciferase, and the normalized values of treated samples were compared to untreated controls.

### Quantitative Real-Time PCR

HEK293T cells were seeded on 6-well dishes for 24 h, then transfected with TOP-Rluc and Firefly Luciferase plasmids for 24 h. Cells were treated with isoproterenol for 6 h in serum-free media. Total RNA was extracted from samples using Quick-RNA Mini-Prep Kit (Zymo, Cat #11-328). Reverse transcription was carried out on 1000 ng purified total RNA with SuperScript II Reverse Transcriptase (Thermo, Cat #18064071). Power SYBR Green (Applied Biosystems, Cat #4368706) and primers described below were used to determine expression of target genes. All *Rluc* expression levels were normalized to housekeeping gene, *GAPDH*.

*Rluc* Forward: TCATGGCCTCGTGAAATCCCGT

*Rluc* Reverse: GCATTGGAAAAGAATCCTGGGTCCG

*GAPDH* Forward: CAATGACCCCTTCATTGACC

*GAPDH* Reverse: GACAAGCTTCCCGTTCTCAG

### β2-AR Internalization Assay

HEK293 cells with stable β2-AR were seeded on 12-well plates and grown to 80% confluency. Cells were treated with 30 µM Dyngo-4a or vehicle (DMSO) for 20 min in serum-free DMEM, then treated with 1 µM isoproterenol for 20 min to induce receptor internalization. Cells were washed with PBS, then stained for 1 h at 4°C in PBS with 1:1000 Alexa-647 conjugated M1 mouse anti-FLAG antibody (Invitrogen, Cat #MA1-91878). Cells were moved to flow cytometry (BD FACS Canto2), and 10,000 cells were analyzed per sample. The mean Alexa 647 of the singlet population was used to quantify surface β2-AR. The percent internalized receptors was calculated as 100% - (Iso surface β2-AR)/(Unstimulated surface β2-AR) × 100%.

### Protein Extraction and Western Blot

HEK293 cells were seeded on 6-well plates and grown to 80% confluency. Cells were serum-starved for 24 h, then pre-treated with vehicle (DMSO), 100 nM Torin, or 30 µM Dyngo-4a for 20 min. Cells were then stimulated with 1 µM Iso for 10 min and lysed with RIPA buffer (Sigma, Cat #R0278) containing protease inhibitors cocktail (Sigma, Cat #P8340), phosphatase inhibitors cocktail (Sigma, Cat #P0044), and 0.1 μM PMSF (Sigma, Cat #P7626). Lysates were then transferred to microcentrifuge tubes and spun for 15 min at 14,000 RPM at 4 °C and the supernatant was used for Western blot experiments. Protein concentration was determined by Pierce BSA Protein Assay Kit (Thermo Fisher, Cat #23225). Protein samples were prepared for western blot analysis by adding 4x sample buffer (Bio-Rad, Cat. #1610747) and 1 mM 2-mercaptoethanol (Sigma, Cat #M6250). Samples were run on a 4-20% MINI-PROTEAN TGX Stain-Free gels (Bio-Rad, Cat #4568093) and transferred to a nitrocellulose membrane for 2 h at 85 V in 4 °C. Membranes were blocked with Intercept Blocking Buffer (LICOR Biosciences, Cat #927-80001) for 1 h and incubated with primary antibodies/TBST against proteins of interest for 3 h at room temperature. Afterwords, membranes were washed, incubated with secondary antibodies for 1 h, and visualized using the Odyssey imager system (LICOR). Bands were quantified and analyzed using ImageStudio software.

## Notes

### Competing Interest Statement

The authors have declared no competing interest.

