## Supplementary Figures for "Extensive location bias of the GPCR-dependent translatome via site-selective activation of mTOR"

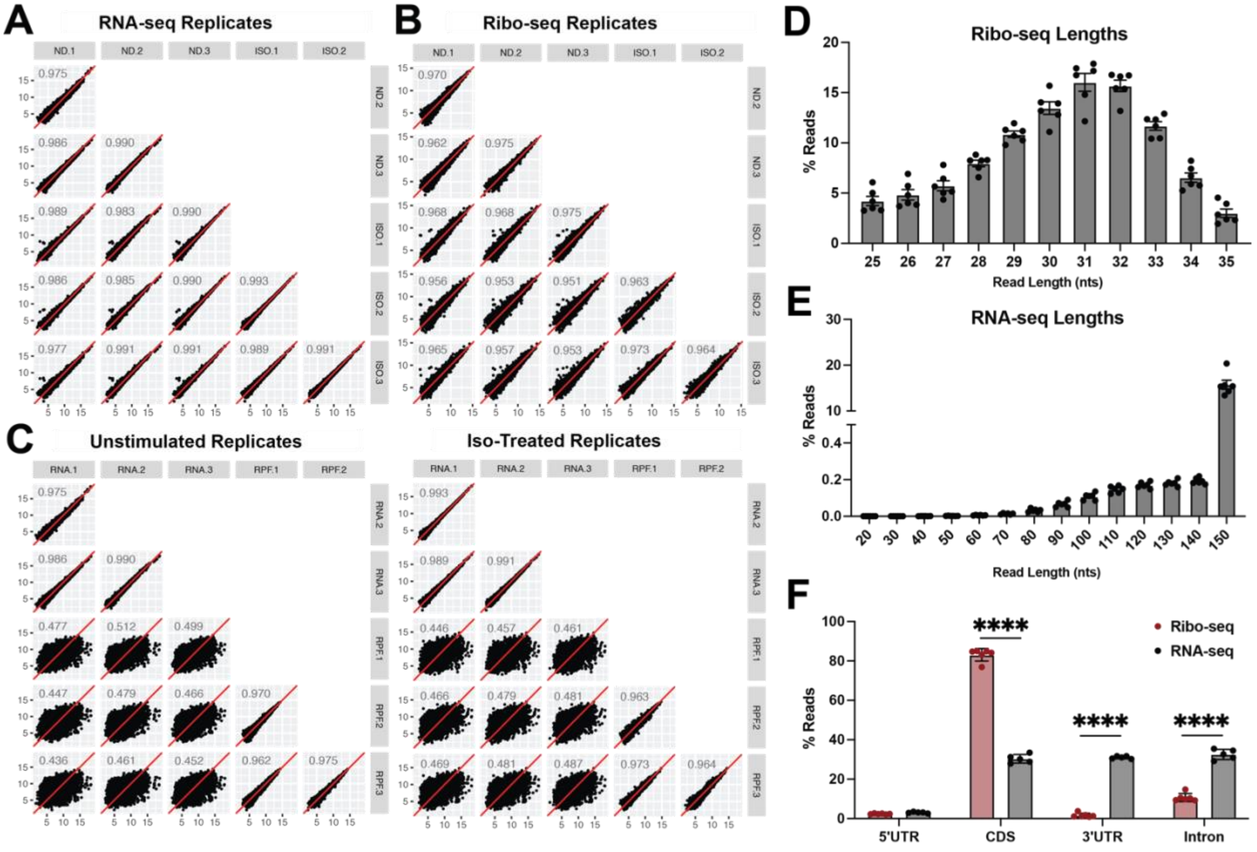


**Figure S1: Ribo-seq quality controls.**

(A-B) Scatter plots depicting the correlation between biological replicates of the RNA-seq (A), Ribo-seq (B), and within-treatment comparisons of unstimulated (left) and Isoproterenol ("Iso")-treated (right) RNA-seq vs Ribo-seq (C). Correlations were calculated with BigPint package in R (Rutter and Cook 2020), and R-squared values are shown. (D) Read length distribution of Ribo-seq samples was calculated using Ribotoolkit (Liu et al. 2020). (E) Read length distribution of the RNA-seq samples was calculated using samtools (Li et al. 2009). (F) Genomic feature type of RNA-seq and Ribo-seq reads were mapped using Ribotoolkit. All bar graphs show mean ± s.e.m. of all n=6 replicates per assay type. **** = $p < 0.0001$ by two-way ANOVA test with Sidak comparing the RNAseq and Ribo-seq experiments.

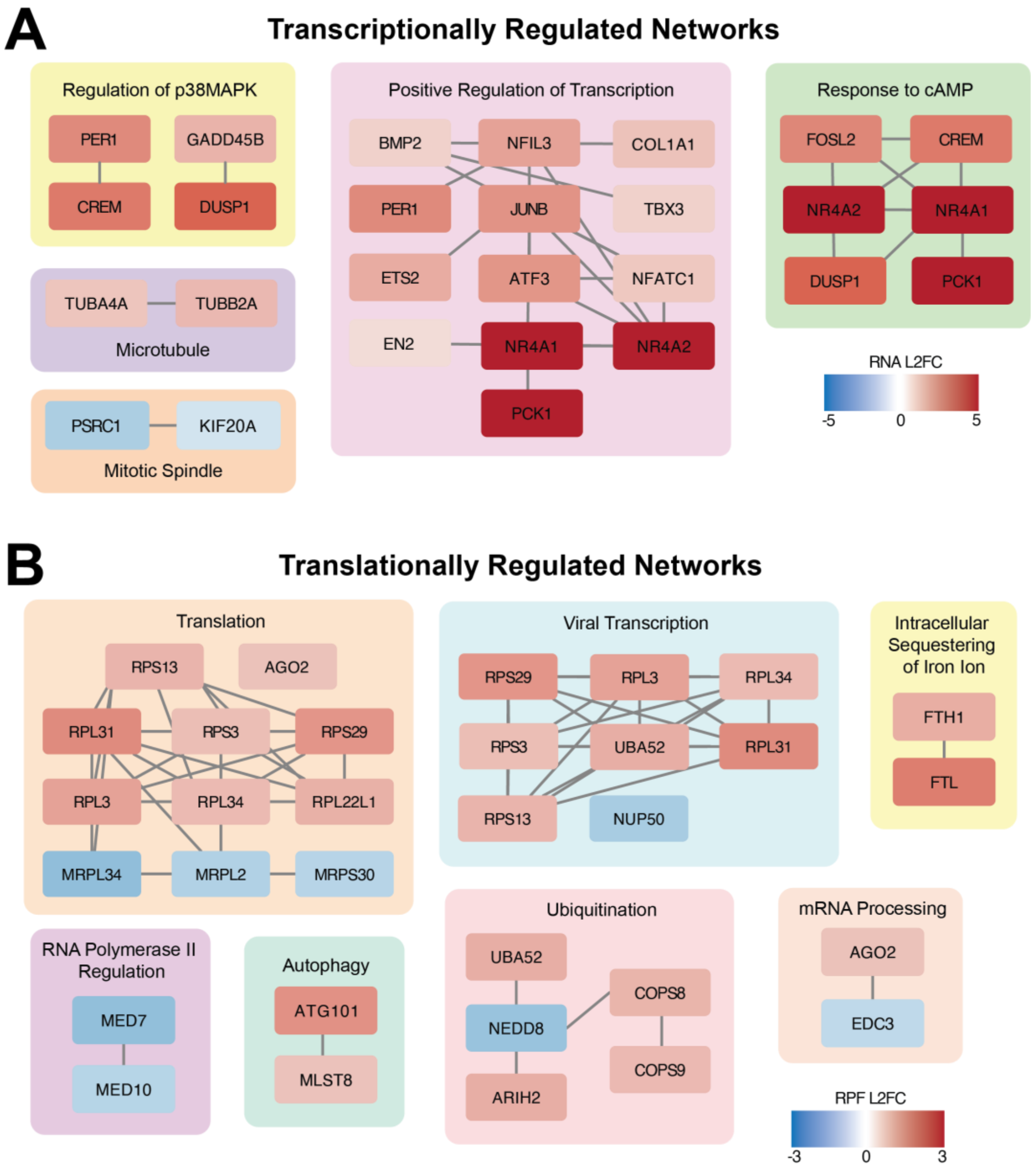


**Figure S2: Transcriptionally and translationally regulated networks by β2-AR activation.**

Transcriptionally regulated genes (A) and translationally regulated genes (B) were analyzed using String-DataBase for network interactions. Resulting networks were edited in Cytoscape (Shannon et al. 2003) and colors of nodes were added to represent RNA-seq (A) or Ribo-seq (B) Log2 fold-change (L2FC).

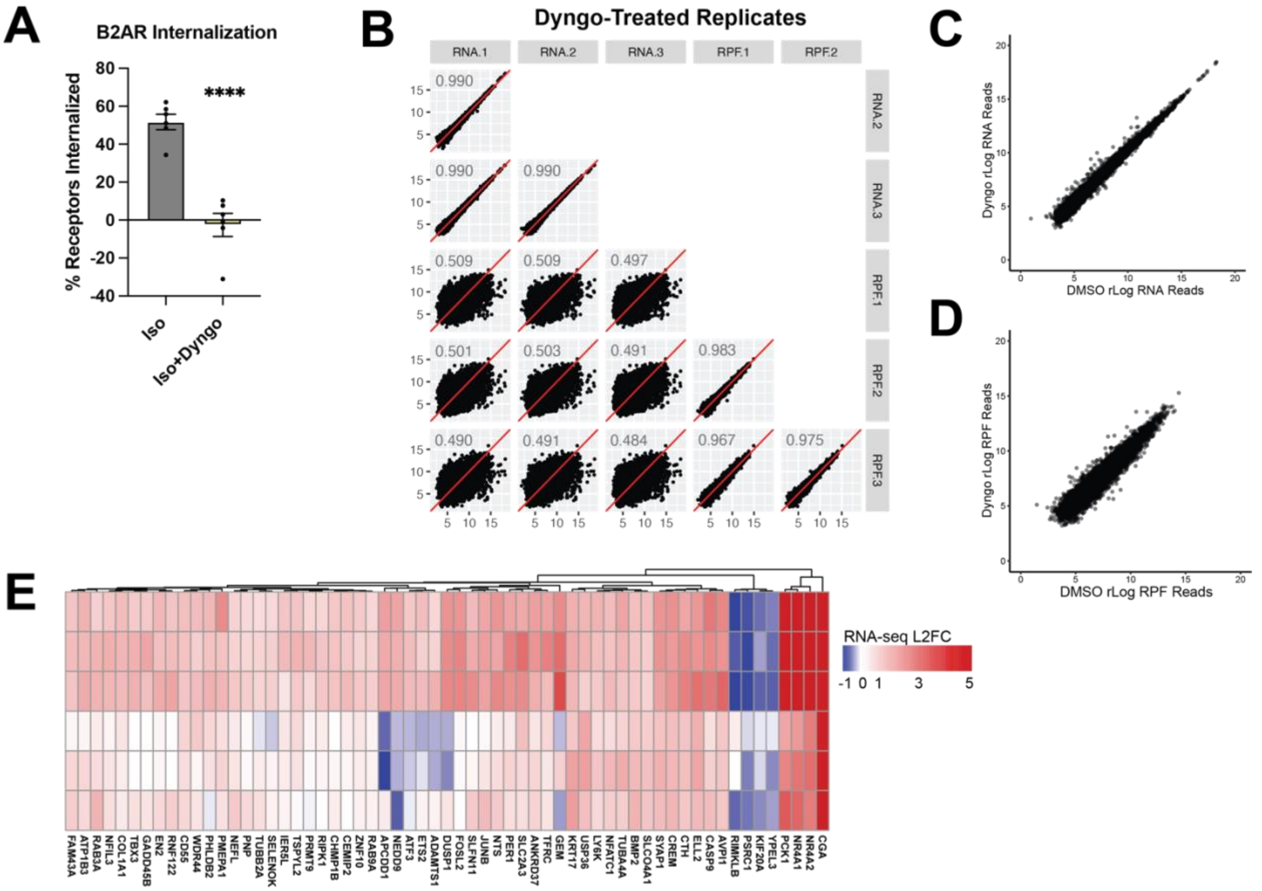


**Figure S3: Dissection of the functional role of β2-ARs endocytosis.**

(A) Isoproterenol ("Iso")-induced β2-AR internalization is inhibited by Dyngo-4a. HEK293 cells stably expressing flag-tagged β2-AR were treated with vehicle (DMSO) or 30 µM Dyngo-4a for 20 min in serum-free medium. Cells were then stimulated with 1 µM Iso for 20 min to induce receptor internalization and analyzed by flow cytometry for surface β2-AR expression. Receptor internalization is calculated as % internalized receptors = 100% - (# surface receptors after 20 min Iso) / (# surface receptors in unstimulated) × 100%. (B) Scatter plots depicting correlation between individual replicates of the Dyngo-4a-treated RNA-seq and Ribo-seq experiments. Correlation was calculated using BigPint package in R (Rutter and Cook 2020). (C-D) Average regularized log reads (rLog calculated in DESeq2 (Love, Huber, and Anders 2014)) are compared between vehicle (DMSO)-treated RNA-seq (C) and Ribo-seq (D) samples. (E) Heatmap of the 61 transcriptional hits from Figure 1. Colors represent Log2 fold-change (Iso-treated vs unstimulated RNA). K-means Clustering was performed with the R-package pheatmap. **** = $p < 0.0001$ by unpaired Student's *t*-test.

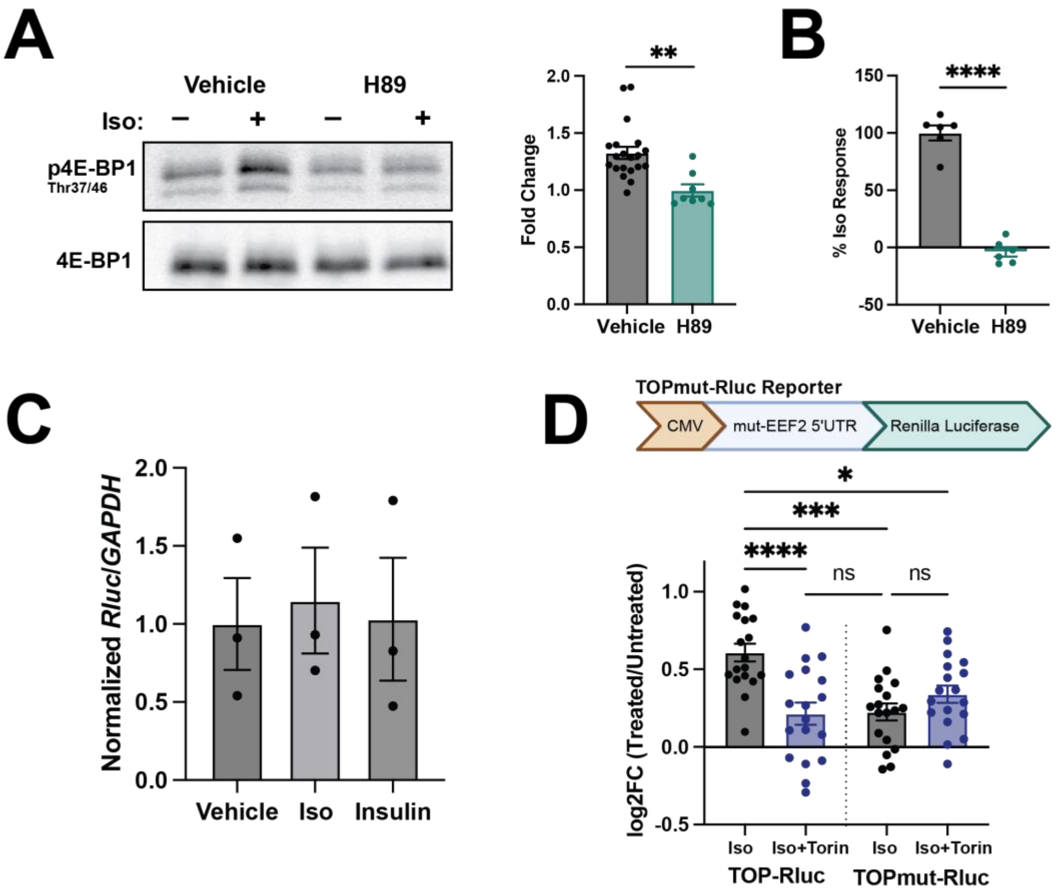


**Figure S4: β2-AR signaling via PKA regulates mTOR activation and translation of TOP mRNAs.**

(A) β2-AR-dependent activation of mTOR requires PKA. HEK293 cells were grown in serum-free DMEM for 24 h, pre-treated with vehicle (DMSO) or 10 µM H-89 dihydrochloride (H89) for 20 min, then treated with 1 µM Isoproterenol (“Iso”) for 10 min. The lysates were analyzed by Western blotting using antibodies against total and phosphorylated 4E-BP1. Fold-change values represent the ratio of phosphorylated/total protein in Iso-treated vs unstimulated conditions. Data are mean of n=8-20 biological replicates. (B) β2-AR-dependent activation of TOP-Rluc translation is mediated by PKA. HEK293T cells transfected with TOP-Rluc and control-Fluc were pre-treated with vehicle (DMSO) or 10 µM H89 for 20 min, then stimulated with 1 µM Iso for 6 h. All values are shown as percentage of the maximum response induced by Iso in the presence of vehicle. Data are mean of n=6 biological replicates. (C) TOP-Rluc mRNA levels do not change following Iso or insulin stimulation. HEK293T cells were transfected with TOP-Rluc and control-FLuc reporters for 24 h, then treated with 100 nM Insulin or 1 µM Iso for 6 h. Cells were lysed and mRNA was analyzed by RT-qPCR using primers for *Renilla luciferase* or *GAPDH*. *Rluc* expression was normalized to *GAPDH*. Data are mean of n=3 biological replicates. (D) Minimal and mTOR-independent induction of TOPmut-Rluc by β2-AR. HEK293T cells transfected with TOPmut-Rluc and control-Fluc were pre-treated with DMSO or 100 nM Torin for 20 min before treatment with 1 µM Iso for 6 h. L2FC is shown relative to the unstimulated condition. Data are mean of n=18 biological replicates. Error bars = ± s.e.m. **** = $p < 0.0001$, ** = $p < 0.01$, * = $p < 0.05$ by unpaired Student’s *t*-test in (A-B) and one-way ANOVA with Dunnett in (C).
